# Effective Clustering for Single Cell Sequencing Cancer Data

**DOI:** 10.1101/586545

**Authors:** Simone Ciccolella, Murray Patterson, Paola Bonizzoni, Gianluca Della Vedova

## Abstract

Single cell sequencing (SCS) technologies provide a level of resolution that makes it indispensable for inferring from a sequenced tumor, evolutionary trees or phylogenies representing an accumulation of cancerous mutations. A drawback of SCS is elevated false negative and missing value rates, resulting in a large space of possible solutions, which in turn makes it difficult, sometimes infeasible using current approaches and tools. One possible solution is to reduce the size of an SCS instance — usually represented as a matrix of presence, absence, and uncertainty of the mutations found in the different sequenced cells — and to infer the tree from this reduced-size instance. In this work, we present a new clustering procedure aimed at clustering such *categorical* vector, or matrix data — here representing SCS instances, called *celluloid*. We show that celluloid clusters mutations with high precision: never pairing too many mutations that are unrelated in the ground truth, but also obtains accurate results in terms of the phylogeny inferred downstream from the reduced instance produced by this method. We demonstrate the usefulness of a clustering step by applying the entire pipeline (clustering + inference method) to a real dataset, showing a significant reduction in the runtime, raising considerably the upper bound on the size of SCS instances which can be solved in practice. Our approach, celluloid: *clustering single cell sequencing data around centroids* is available at https://github.com/AlgoLab/celluloid/ under an MIT license, as well as on the *Python Package Index* (PyPI) at https://pypi.org/project/celluloid-clust/

## I. Introduction

A tumor, at the time of detection, usually by performing a biopsy on the extracted tissue, is the result of a tumultuous evolutionary process, originating from a single tumor cell — the founder cell [30] — that has acquired a set of *driver* mutations, which inhibits control on the proliferation of subsequent cancer cells. From that moment, the combination of unrestrained proliferation and a very hostile environment — as the immune system fights for survival, the tumor cells, under extreme selection pressure, have to disguise themselves to avoid being attacked, compete with each other, all while having to thrive under low levels of oxygen — produces the accumulation of a highly elevated number of mutations, including structural variations. This model of tumor evolution is called the clonal model [30], since a clone is a population of cells carrying the same set of mutations. Understanding this clonal evolutionary history can help clinicians understand the cell heterogeneity in the tumors of various types of cancer and, more importantly, gives insights on how to devise therapeutic strategies [28], [38].

The rise of *next-generation sequencing* (NGS) technologies has led to the computational problem of tumor phylogeny inference from NGS bulk sequencing data [8], [12], [23], [35]. This idea is very cost-effective, since NGS data is cheap to obtain and very reliable. The procedure consists of extracting different samples of the tumor and aligning the NGS reads against the reference genome: this allows to determine the approximate fraction of reads from each sample that are affected by any given mutation. This fraction is taken as a proxy of the fraction of cells in each sample that are affected by that mutation. The main difficulties of this technique are the fact that a sample contains a mix of both healthy cells and cancer cells, while the cancer cells are an unknown mixture of different clones.

Since both the composition of the samples and the evolutionary history are unknown in this model, most of the approaches relied on some simplifying assumption, such as the *infinite sites assumption* (ISA) which postulates that each mutation is acquired exactly once, and never lost, in the entire tree. While this assumption has only limited biological validity [3], [22], it reduces greatly the space of all possible solutions, making feasible several approaches [8], [12], which are at least partially inspired by a classical linear time algorithm [11] for reconstructing the phylogeny from noise-free character data. This latter computational problem is called the *perfect phylogeny* problem.

The newer *single cell sequencing* (SCS) technology provides a much finer level of resolution: in fact we can determine whether or not a given cell *has* a mutation, therefore avoiding the notion of sample and the approximations implied by the use of samples. Still, SCS is expensive and plagued by high dropout, *i.e.*, missing values, and false negative rates.

Nowadays, we are witnessing a decrease in SCS costs, coupled with improvements in the dropout and false negative rates, stimulating the research on tools for tumor evolution inference from SCS data [5]–[7], [17], [32], [39]. We believe that this line of research is going to become even more important in the next few years, since currently available SCS data is associated with a very large solution space. Moreover, the high missing value and false negative rates allow a huge number of possible phylogenies with near optimal values of the objective function. This fact makes difficult to determine which methods actually produce better solutions, and shows that the objective function is not able to fully capture the biological soundness of the phylogeny.

These advances in costs and quality of the data produced will result in larger, but more constrained, instances — the net effect being a considerable reduction in the number of likely solutions. Since most of the currently available methods do not scale well to large instances (usually their running time is quadratic with respect to the number of mutations), one solution could be to reduce the size of an instance by clustering: for example, SPhyR [7] uses *k*-means [1], [25] to such purpose. However, *k*-means is designed for continuous data — where means are usually based on a Euclidean distance — while SCS data, specifying the presence (1) or absence (0) of a mutation in a cell, or the fact that it is missing (2), can be thought of as categorical.

Clustering categorical data is an active field of research in data mining [19], where massive databases of categorical data are handled, *e.g.*, finding groups of members of a particular insurance policy who also travel overseas on a regular basis. While the goal of clustering here is to allow faster downstream phylogeny inference, it could also be used for error correction in SCS data [26], [29] — a closely related topic.

In this paper we devise a new method for clustering categorical data, called *celluloid* — a schematic of this depicted in Figure 1. Not only does *celluloid* cluster categorical data (based on a *k*-modes framework), but because of its novel *conflict dissimilarity* measure — tailored to properties specific to SCS data (see Section II: Methods) — it obtains the best results in a comparison of a variety of different clustering approaches on SCS data. This comparison has three goals: (1) to assess which clustering methods are more precise on SCS data by comparing the actual clustering produced, (2) to evaluate the effect of a clustering step on the downstream phylogeny inference step, and (3) to assess the usefulness of a clustering step on real data, by showing its capability to reduce an instance that is too large for some of the current methods, so that the clustered instance can be easily solved.

**Fig. 1.**
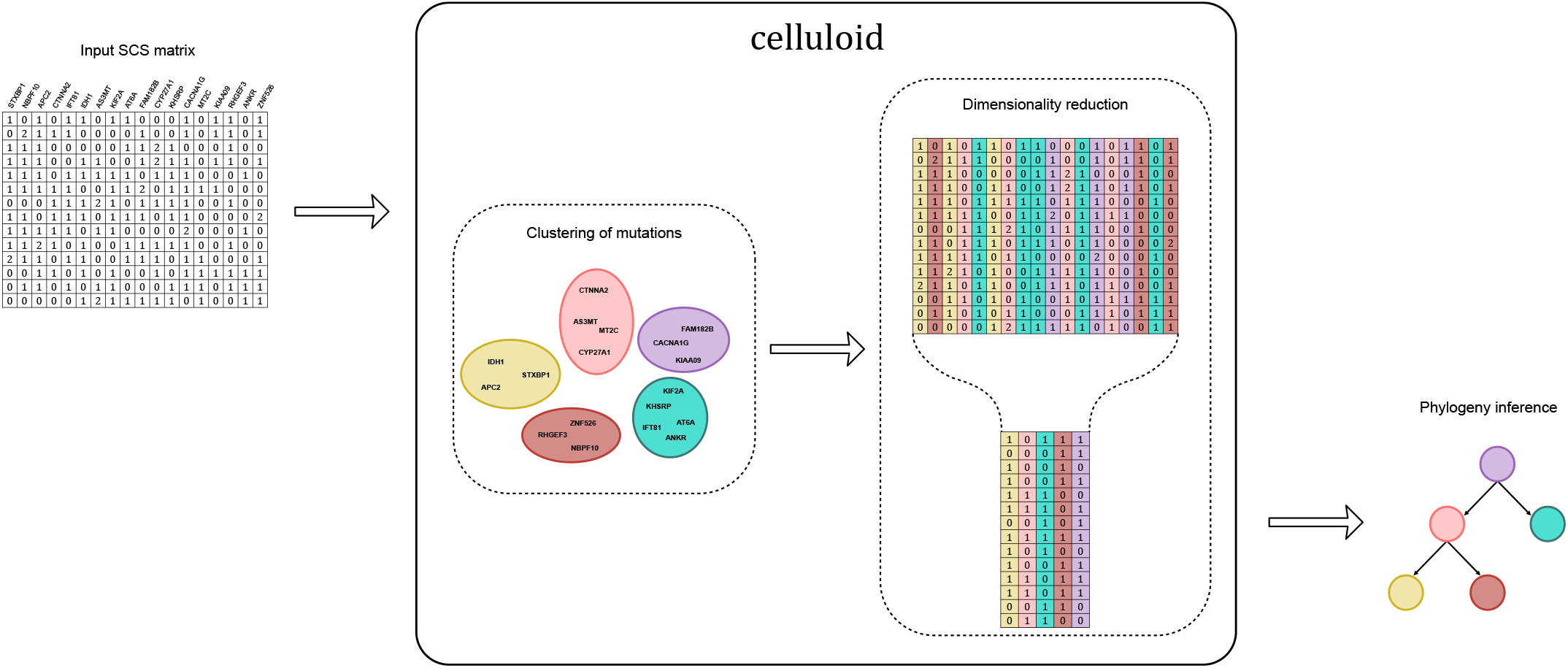
A schematic overview of how *celluloid* works, and how it fits into a cancer phylogeny inference pipeline. When an input SCS matrix (upper left) is too large for a phylogeny inference method (lower right), one can use *celluloid* (middle) as an intermediate step to reduce the dimensionality of this instance, making feasible the phylogeny inference.

## II. Methods

Starting with the method we devise, *celluloid*, we now give an overview of the methods we consider for reducing the size of single-cell sequencing (SCS) datasets, by clustering mutations. Specifically, the problem is: given *m* objects, each with *n* values — each value taken from a set of possible categories — we wish to cluster the *m* objects into *k* groups. In this context, the objects are *m* mutations (columns of an SCS matrix), each one over *n* cells (rows of an SCS matrix), while each value can be one of three possibilities — that the mutation is *present* (1), *absent* (0), or *missing* (2) from the cell.

Again, the goal of clustering here is to allow faster downstream phylogeny inference. This is why we devise a method which takes, *a priori*, parameter *k*: the desired number of clusters, i.e., the largest number that the downstream inference can complete in a reasonable amount of time. For the same reason, we also restrict our comparison to such methods, for which there is already a wide variety. In light of this, we did not consider the entire field of methods which determine the number of clusters based on properties of the data [2], [20], [27], which could also be appropriate.

### A. Celluloid

Clustering methods are based on a notion of distance (or of similarity) between the elements that we want to cluster. In our case, we are dealing with a set *X* of *m* objects on *n categorical attributes* {*A*_1_…, *A_n_*} — in other words, each object *x* ∈ *X* is a point 〈*x*[1],…, *x*[*n*]〉 of the *n*-dimensional space 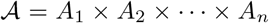 — where an object *x* has a *categorical value x*[*i*] ∈ *A_i_* for each attribute *A_i_*.

An intuitive and widely used notion of distance is the the *dissimilarity measure d_M_*, also called *matching dissimilarity*, between two objects *x* = 〈*x*[1],…, *x*[*n*]〉 and *y* = 〈*y* [1],…, *y*[*n*]〉 of 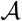, defined as:

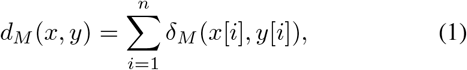

where *δ_M_* is a trivially defined dissimilarity, based on the identity of two categorical values:

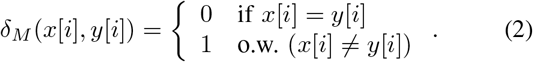

In our context, the value 2 is special, since it represents *missing* data. Therefore, the fact that a given value is equal to 2 should not be penalized. For this reason, we design a new dissimilarity, based on the notion of *conflict* (Equation 4) by adapting the previous matching dissimilarity of Equation 2 to obtain what we call the *conflict dissimilarity*, defined as:

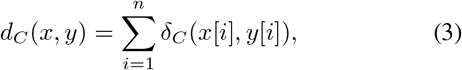

and *δ_C_* is a slight relaxation of *δ_M_*, where 2s are not penalized:

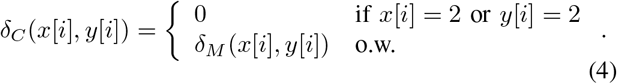

Our method *celluloid* is based on the *k*-modes framework [14], [15] which clusters *m* objects *X* on 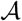, by computing an *m* × *k partition matrix* [13] *p*[·, ·] — a partitioning of these *m* objects into *k* clusters — and a set *Q* = {*q*_1_,…, *q_k_*} of *k modes* and that minimize the following objective function:

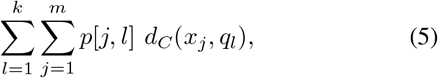

subject to

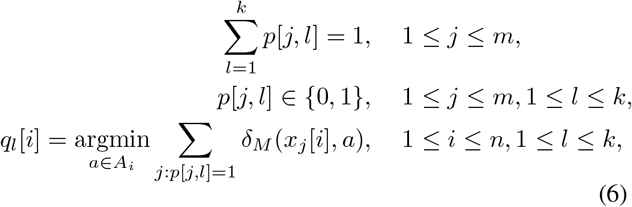

where *p*[*j, l*] = 1 if and only if the object *x_j_* is placed in the cluster *C_l_* whose mode is *q_l_*.

That is, for each *q_l_* ∈ *Q, q_l_*[*i*] is one of the possible categorical values of the attribute *A_i_*. Most precisely, *q_l_*[*i*] is a *mode* among the values (of the attribute *A_i_*) of the objects that are in the cluster *C_l_*. Note that every *A_i_* = {0,1,2} when considering single cell sequencing data for inferring tumor evolution.

Now, a *mode* of cluster *C_l_* ⊆ *X* is a vector *q_l_* which minimizes:

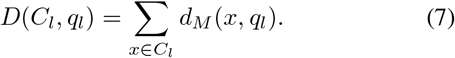

Note that *q_l_* is not necessarily an element of *C_l_*. The *k*-modes algorithm then starts with some initial set *Q*^0^ of *k* modes, and an initial collection 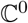 of *k* disjoint subsets of *X*, and then iterates the operations:

1. compute *d_C_*(*x, q_l_*), where *q_l_* is a mode of cluster 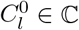, for each object *x* and each cluster 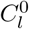;
2. for each mode *q_l_* create an empty cluster 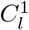;
3. allocate *x* to a cluster 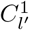 minimizing *d_C_*(*x, q_l_*,), hence updating *p*[·, ·]; and
4. recompute a new set of modes *Q*^1^ according to the new clusters 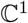, exploiting the last constraint of Equation 6;

until convergence, i.e., until *Q*^*t*+1^ = *Q^t^*. The modes in the third step above are found according to the following theorem, where *c_r_*[*i*] is the *r*-th categorical value of the attribute *A_i_* and *f*(*A_i_* = *c_r_*[*i*]|*X*) is its relative frequency in *X*.

#### Theorem 1 ([15])

Eq. 7 is minimized iff *f*(*A_i_* = *q_l_*[*i*]|*X*) ≥ *f*(*A_i_* = *c_r_*[*i*]|*X*) for *q_l_*[*i*] ≠ *c_r_*[*i*] ∀ *i* ∈ [1…*n*]. In other words, Eq. 7 is minimized by selecting any mode value for each attribute. Note that this theorem implies that the mode of *X* is not necessarily unique.

Since the solution depends on the initial set *Q*^0^ of *k* modes, we consider two procedures for initializing *Q*^0^. The first one is quite simple: a random selection of *k* objects from the set *X* of objects as the initial *k* modes — which we refer to as the *random initialization* procedure. The second one, devised in [15], is a more complicated procedure, based on the frequencies *f* (*A_i_* = *c_r_*[*i*]|*X*) of all categories, which we refer to as the *Huang initialization* procedure [15]. We focus on this second procedure, since it achieves the best results:

1. order the categories of each attribute *A_i_* in descending order of frequency, i.e., *f*(*c*_*r*_1__[*i*]) ≥ *f*(*c*_*r*_2__ [*i*]) ≥ *f*(*c*_*r*_3__[*i*]), *etc*.;
2. assign uniformly at random the most frequent categories to the initial *k* modes;
3. for each mode *q_l_* obtained in the previous step, select the *x_j_* ∈ *X* most similar to *q_l_* and make this *x_j_* the mode 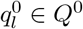, such that 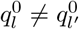 for *l* ≠ *l*′.

This final step is to avoid empty clusters in 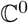. This initial selection procedure is aimed at having a diverse set of initial modes *Q*^0^, which can lead to better clustering results — see more details in [15].

Our approach, *celluloid: clustering single cell sequencing data around centroids*, is our conflict dissimilarity and the Huang initialization procedure, used within the *k*-modes framework — its implementation available at under an MIT license [43]. It is also available on the *Python Package Index* (PyPI) [44]: installable via `pip`, *e.g.*, `pip install celluloid-clust`.

### B. k-*modes*

Since there is package for computing a clustering based on the *k*-modes framework available on the *Python Package Index* (PyPI) [45], we decided to use this in our tests as well. The above package comes with a variety of standard dissimilarity measures, as well as the Huang (and random) initialization procedures (see above).

Here we use the above implementation with the options of matching dissimilarity (Equation 2) and the Huang initialization procedure, since this combination of options produced the best results. This is what we refer to as *k-modes* in our experiments.

### C. k-*means*

Given *m* objects *X* = {*x*_1_,…,*x_m_*} on *n real* values, *i.e.*, each *x_i_* ∈ ℝ^*n*^, and an integer *k*, the *k*-means algorithm [1], [25] finds the vector of *k* values, called *means, Q* = {*q*_1_,…,*q_k_*} and an *m* × *k* partition matrix minimizing the following objective function:

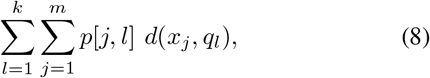

subject to the first two constraints in Equation 6, while each *mean q_l_* is based on a *distance measure d*, that is usually the Euclidean distance, *i.e*.,

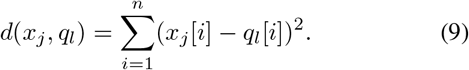

The *k*-means algorithm starts with some initial set *Q*^0^ of *k* means, and an initial collection 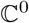 of *k* disjoint subsets of *X*, and then iterates the operations 1–3 as in the *k*-modes algorithm, but using the Euclidean distance (Equation 9) instead of dissimilarity, and computing *means* instead of modes, that minimize instead the objective function of Equation 8, subject to the first two constraints in Equation 6.

In our experiments, we used the implementation of *k*-means clustering available in scikit-learn [46].

### D. Affinity propagation

The affinity propagation algorithm [10] uses as input a set of similarities between data points. Those similarities are clustered by choosing a point as a representative for each class. Such points gradually emerge iteratively using a message-passing procedure, where each point exchanges messages with all other points.

In particular, there exist two types of messages: (1) *responsibility r*(*i, k*) is sent from point *i* to the candidate representative point *k* reflecting the cumulative evidence of how well-suited *k* is to serve as representative of *i*; and (2) *availability a*(*i, k*) sent from the candidate representative *k* to *i* reflecting the cumulative evidence of how well-suited *k* should be chosen by *i* to be its representative. Both variables take into account other potential candidates.

Such messages are exchanged iteratively, each time refining *r*(·,·) and *a*(·,·), until a stop condition is fulfilled. At any point these variables can be combined to identify the representatives. For each point *i*, the value of *k* that maximizes *r*(*i, k*) + *a*(*i, k*) identifies the representative *k* of *i*, where *k* and *i* can be the same point.

In our experiments, we used the implementation of affinity propagation clustering available in scikit-learn [47].

### E. Hierarchical agglomerative clustering

The hierarchical agglomerative clustering method [18] produces, in an hierarchical fashion, groups of disjoint subsets of a given set, each maximizing an internal similarity score. The procedure is executed in a bottom-up fashion in which, initially, each point is in a subset by itself. At each step, two sets are merged together, so that the union maximizes a given criterion. The procedure is then repeated until only one group remains, thus having the complete hierarchical structure of the clustering.

In our experiments we used the Manhattan metric, *i.e*., the sum of the horizontal and vertical distance between two points, to compute the similarity scores, as this metric is more suited for a matrix containing categorical data, as opposed to the Euclidean distance more suited for a continuous-valued matrix. The similarity of two sets of observations is computed using the average of the distances between elements in the respective sets. Consequently, as expected, the hierarchical agglomerative clustering, when using the Manhattan metric, performed better (see Section III: Results) than the Euclidean metric, and so we only included the former in the comparison.

In our experiments, we used the implementation of agglomerative hierarchical clustering available in scikit-learn [48].

### F. BIRCH clustering

The BIRCH clustering procedure [40] takes as input a set of points as well as the desired number of clusters, and operates in four steps. (1) In the first step it computes the *clustering feature* (CF) tree while computing measures using a predefined metric. (2) The second optional phase tries to build a smaller CF tree while removing outliers and regrouping crowded subclusters into larger ones. (3) In the third step, a hierarchical clustering algorithm, usually an adaptation of the agglomerative clustering (briefly described in the previous subsection), is used to cluster all the leaves of the CF tree. During this phase, the centroids of each cluster are computed. (4) The centroids are then used for the final fourth step to further refine the clusters, since minor and localized misplacements could occur during the previous step. This last step can also be used to label the points with the cluster they are placed in — we used this feature to obtain the groups of mutations in our experiments.

In our experiments, we used the implementation of BIRCH clustering available in scikit-learn [49].

### G. Spectral clustering

The spectral clustering algorithm [34] — see [37] for a gentle and comprehensive treatment of the topic — first performs dimensionality reduction on the data, and then clusters, using a standard clustering technique such as *k*-means, the data in this reduced dimension. In order to reduce the dimensionality, it first constructs a similarity *graph* from the initial matrix of similarities of the input set of data points — usually a sparse representation of this similarity matrix, *e.g*., the *k*-nearest neighbor graph [37]. It then takes the Laplacian matrix [4] on this graph, and a subset of the (relevant) eigenvectors (spectrum) of this Laplacian matrix — itself a matrix, is the reduced-dimension data. It is then this matrix which is passed to the clustering technique.

In our experiments, we used the implementation of spectral clustering available in scikit-learn [50].

### H. Generation of simulated data

The simulated data are generated as follows. First we simulate a random tree topology on s nodes, each representing a tumor clone, by first creating a root (the germline) and then iteratively attaching the *s* – 1 remaining nodes uniformly at random to any other node in the tree. The nodes are then randomly labeled with *m* mutations — meaning that each mutation is acquired at the node that it labels. Then, a total of *n* cells are associated to the nodes, uniformly at random. A binary matrix *M* is then extracted from these cells giving rise to a genotype profile for each cell (a row in *M*), which is the presence (1) or absence (0) of each mutation (column in *M*) in the cell, given the presence or absence of the mutations on the path in the tree from the root to this cell. The binary matrix is then perturbed according to the false negative, false positive and missing value rates, to simulate a real-case scenario. Each of the s nodes is therefore considered as a natural (true) cluster of the simulated dataset.

## III. Results

To evaluate the accuracy of the clustering methods, we designed a two-fold experiment on synthetic datasets. We first measure the quality of the clusters found by the methods, and later we evaluate how such clusters impact the quality of the phylogenies returned by a cancer progression inference method.

For each experiment, we consider 100, 200 and 300 cells, respectively experiments 1, 2 and 3. For each such value we generated 50 simulated datasets (according to Section II-H), where we fixed the number s of clones to 20 and the number *m* of mutations to 1000. While this number of mutations is at the high end in terms of currently available real cases, it will be a typical size in the near future — some such cases already existing today (see Section III-C).

We performed clustering on the datasets of our experiments to obtain instances with a reduced number of columns (mutations), which can in turn be given as input to such a cancer progression inference method above. The clustering methods we used were all the ones described in the Methods section (Section II), *i.e*., *celluloid, k*-modes, *k*-means, affinity, agglomerative, BIRCH and spectral clustering. Note that all such clustering methods are general-purpose: given *m* objects on *n* categorical values, and an integer *k*, each method clusters the *m* objects into *k* groups. Since the cancer inference methods tend to scale quadratically with the number of mutations, we decided to choose a *k* of 100 for each method, which is a reasonable number of mutations given the currently available literature. Note that experiments for additional parameter settings can be found in the Supplementary Material document.

### A. Evaluating a clustering

To evaluate the clusters obtained, we used standard precision and recall measures, adapted to the particular goal, as follows.

**Precision**: measures how well mutations are clustered together. For each pair of mutations appearing in the same clone in the simulated tree, we check if they are in the same cluster, resulting in a *true positive* (*TP*). For each pair of mutations clustered together that are not in the same clone, we encounter a *false positive* (*FP*). The value of the precision is then calculated with the standard formula: 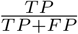.
**Recall**: measures how well mutations are separated. For each pair of mutations in the same clone, we now also check if they are not in the same cluster, resulting in a *false negative* (*FN*). The recall is then calculated as: 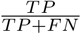.

It is important to highlight that we are mostly interested in obtaining a high *precision* since, while cancer phylogeny inference algorithms can later cluster together mutations by assigning them to same subtree or the same path, they cannot separate mutations that have been erroneously clustered together. It is for this same reason that the *number* (*k*) of clusters we chose is simply the largest such that the downstream inference tool can complete in a reasonable amount of time.

In Figures 2, 3 and 4 — representing the respective experiments 1, 2 and 3 — a common trend is evident: indeed, standard clustering methods (*k*-means, affinity, agglomerative, BIRCH and spectral clustering) perform much more poorly than *celluloid* and *k*-modes, presenting a gap from all the other methods in terms of both precision and recall. On the other hand, *celluloid* and *k*-modes differ slightly in terms of precision.

**Fig. 2.**
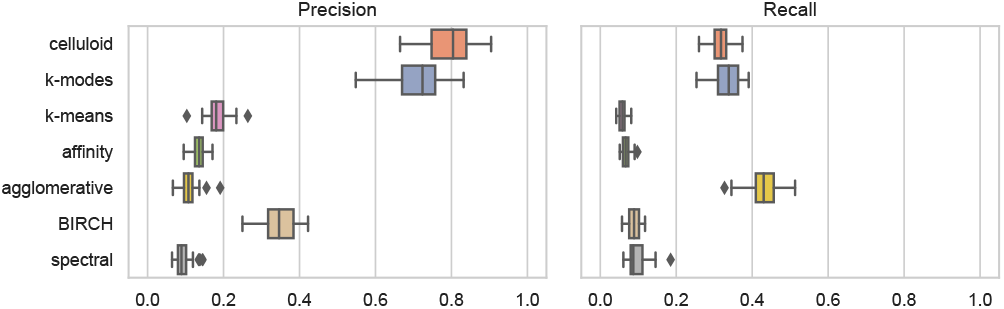
Precision and recall results for experiment 1, generated with a total of 1000 mutations, 100 cells and a clustering size of ***k*** = 100. The plots include results for *celluloid*, ***k***-modes, ***k***-means, affinity, agglomerative, BIRCH and spectral clustering.

**Fig. 3.**
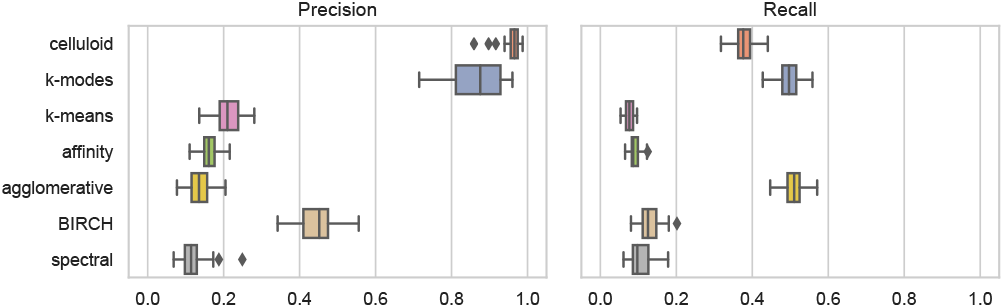
Precision and recall results for experiment 2, generated with a total of 1000 mutations, 200 cells and a clustering size of ***k*** = 100. The plots include results for *celluloid*, ***k***-modes, ***k***-means, affinity, agglomerative, BIRCH and spectral clustering.

**Fig. 4.**
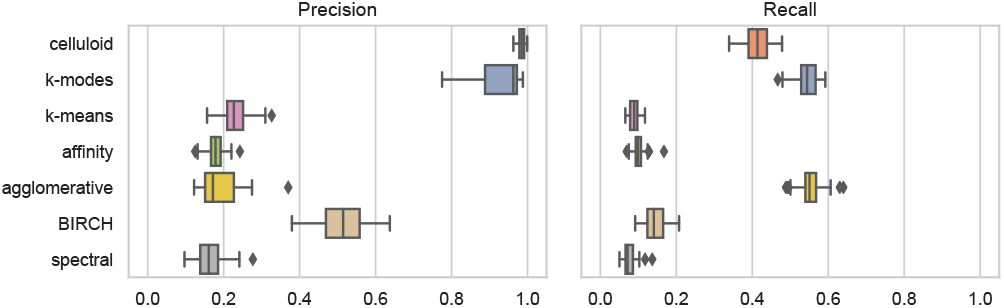
Precision and recall results for experiment 3, generated with a total of 1000 mutations, 300 cells and a clustering size of ***k*** = 100. The plots include results for *celluloid*, ***k***-modes, ***k***-means, affinity, agglomerative, BIRCH and spectral clustering.

It is interesting to notice that the precision of the conflict dissimilarity rapidly increases when the amount of cells increase, thus being well-suited for future increases on the size of SCS datasets.

We have also evaluated the quality of the clusters with several standard clustering metrics such as the adjusted Rand index [16], Fowlkes-Mallows index [9], completeness score [31] and V-measure [31]. In this case, we measure how much the computed clustering is similar to the ground truth.

In Figures 5, 6 and 7 — representing again, respectively, the results of experiments 1, 2 and 3 — we see the similar trend of *celluloid* and *k*-modes performing much better than all the other methods, presenting this same gap from all the other methods, while differing slightly from each other.

**Fig. 5.**
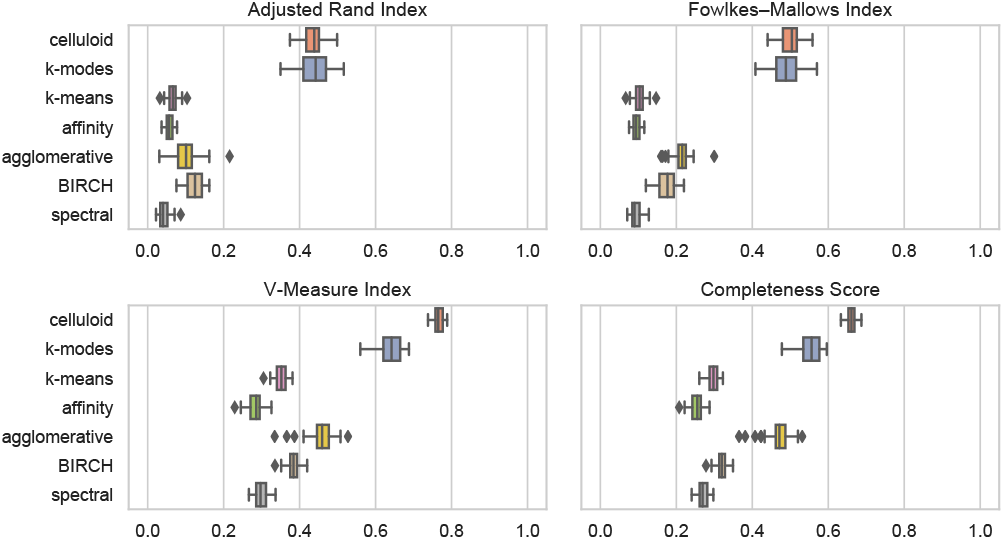
The adjusted Rand index, Fowlkes-Mallows index, completeness score and V-measure between all clustering methods and the ground truth for experiment 1, generated with a total of 1000 mutations, 100 cells and a clustering size of ***k*** = 100. The plots include results for *celluloid*, ***k***-modes, ***k***-means, affinity, agglomerative, BIRCH and spectral clustering.

**Fig. 6.**
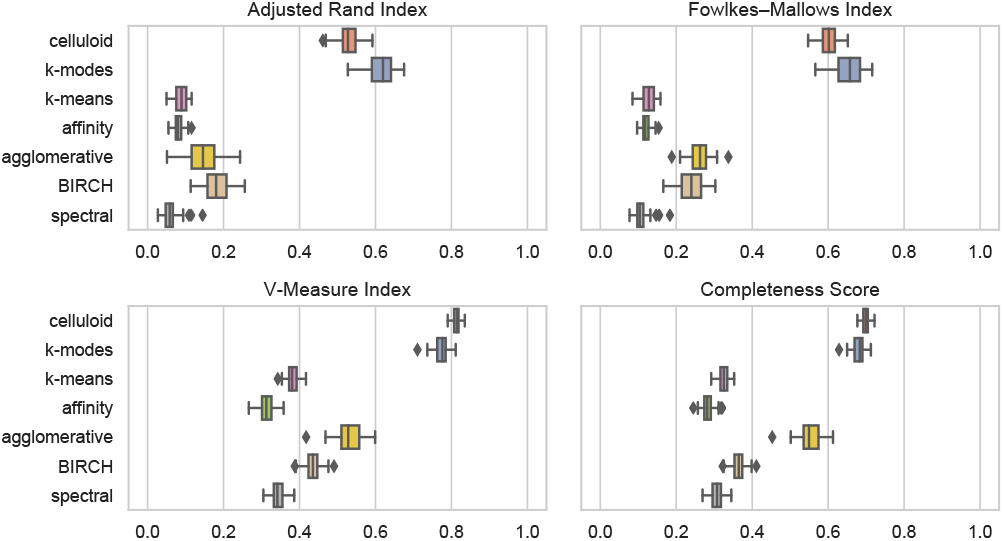
The adjusted Rand index, Fowlkes-Mallows index, completeness score and V-measure between all clustering methods and the ground truth for experiment 2, generated with a total of 1000 mutations, 200 cells and a clustering size of ***k*** = 100. The plots include results for *celluloid*, ***k***-modes, ***k***-means, affinity, agglomerative, BIRCH and spectral clustering.

**Fig. 7.**
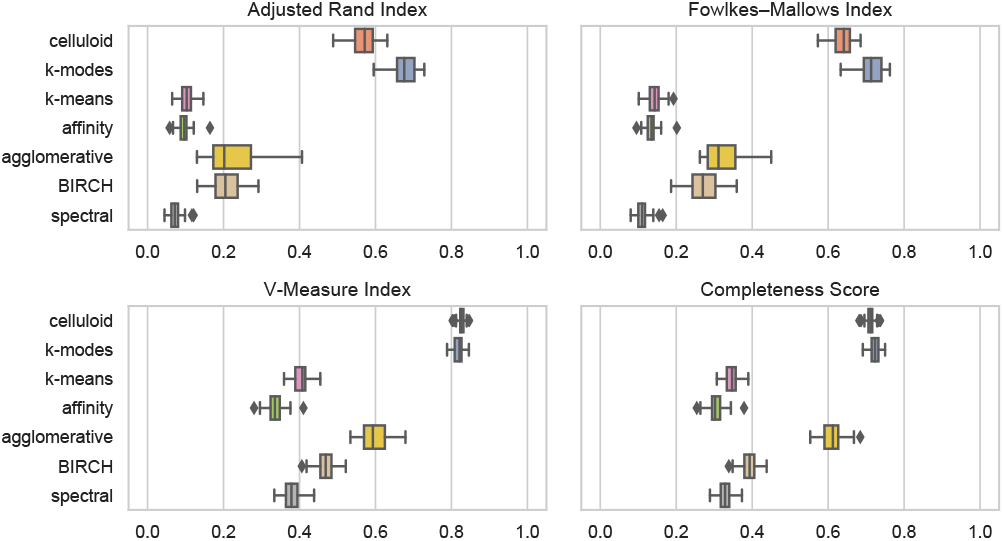
The adjusted Rand index, Fowlkes-Mallows index, completeness score and V-measure between all clustering methods and the ground truth for experiment 3, generated with a total of 1000 mutations, 300 cells and a clustering size of ***k*** = 100. The plots include results for *celluloid*, ***k***-modes, ***k***-means, affinity, agglomerative, BIRCH and spectral clustering.

We decided also to compute the adjusted Rand index, Fowlkes-Mallows index, completeness score and V-measure for *all pairs* of techniques to allow the observation of the similarity, according to the measures, between different algorithms. The heatmaps in figures 8, 9 and 10 — representing, respectively, experiments 1, 2 and 3 — show the average value of the scores of each simulated dataset for each pair of clustering method.

**Fig. 8.**
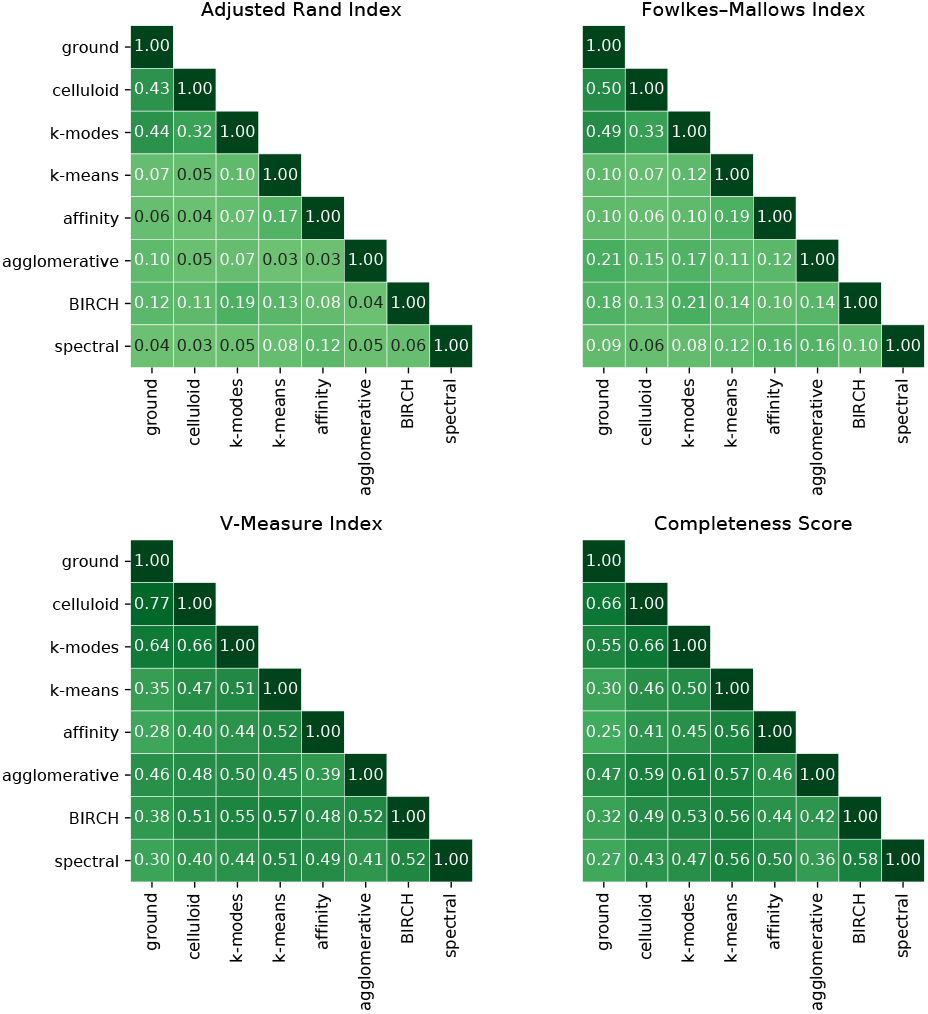
The adjusted Rand index, Fowlkes-Mallows index, completeness score and V-measure between all pairs of clustering methods for experiment 1, generated with a total of 1000 mutations, 100 cells and a clustering size of ***k*** = 100. The plots include results for *celluloid*, ***k***-modes, ***k***-means, affinity, agglomerative, BIRCH and spectral clustering. Each cell is the average of the scores obtained for each simulated instance.

**Fig. 9.**
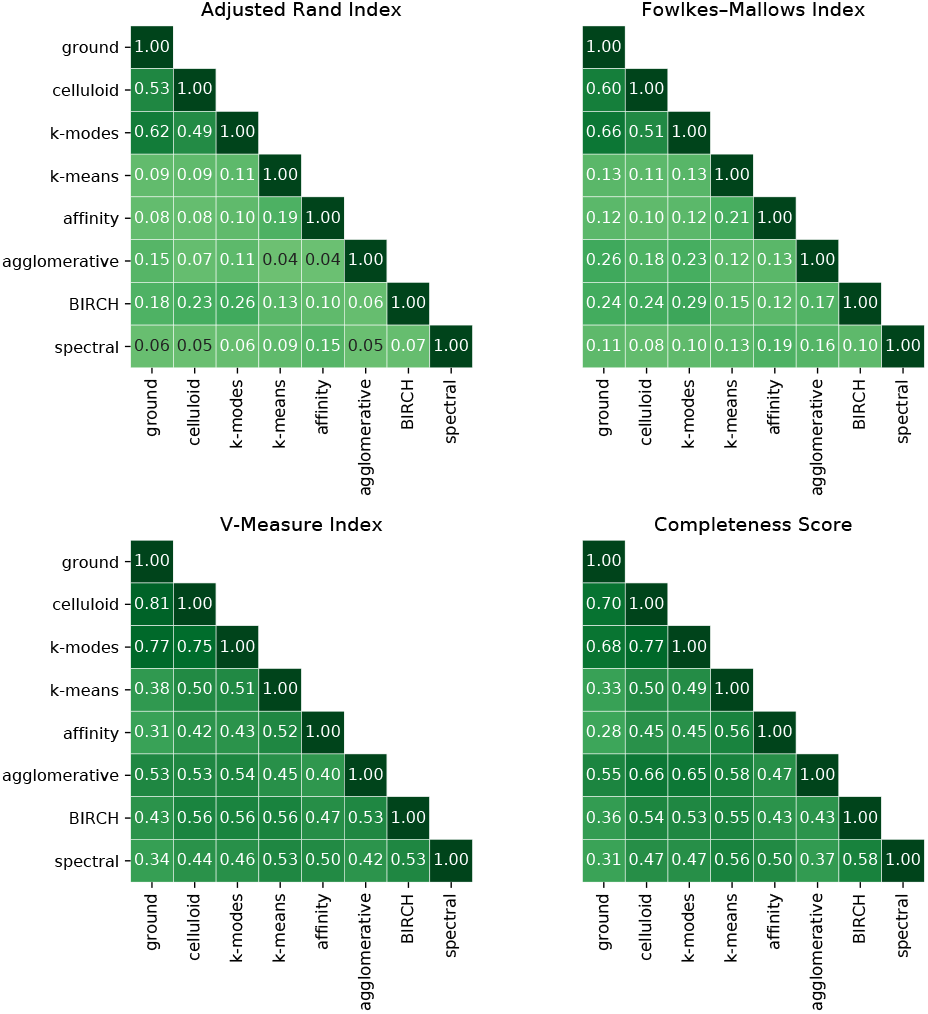
The adjusted Rand index, Fowlkes-Mallows index, completeness score and V-measure between all pairs of clustering methods for experiment 2, generated with a total of 1000 mutations, 200 cells and a clustering size of ***k*** = 100. The plots include results for *celluloid*, ***k***-modes, ***k***-means, affinity, agglomerative, BIRCH and spectral clustering. Each cell is the average of the scores obtained for each simulated instance.

**Fig. 10.**
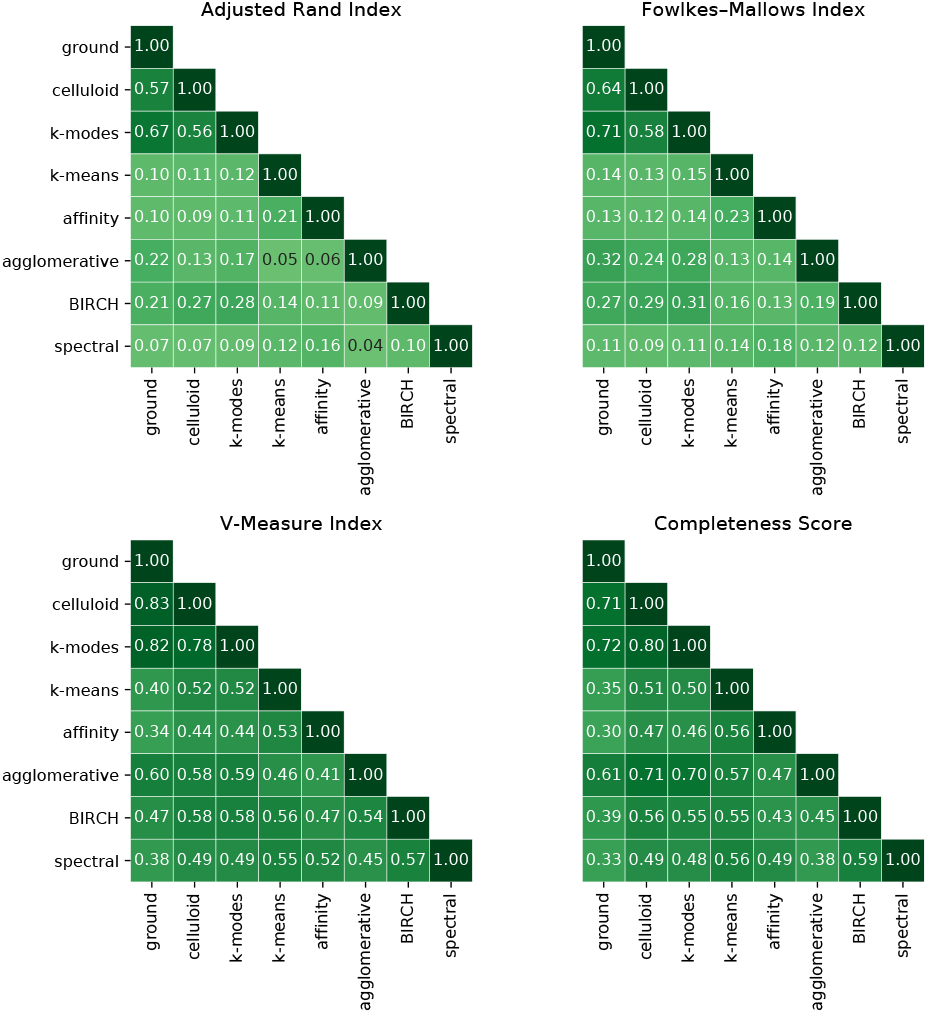
The adjusted Rand index, Fowlkes-Mallows index, completeness score and V-measure between all pairs of clustering methods for experiment 3, generated with a total of 1000 mutations, 300 cells and a clustering size of ***k*** = 100. The plots include results for *celluloid*, ***k***-modes, ***k***-means, affinity, agglomerative, BIRCH and spectral clustering. Each cell is the average of the scores obtained for each simulated instance.

The first column of all the heatmaps represents the comparison with the ground truth, where in all the cases *celluloid* achieves larger values, thus more resembling the ground truth. Furthermore, *celluloid* and *k*-modes are very similar to each other, while the other methods tend to be quite dissimilar to each other.

### B. Assessing the impact of a clustering

To better understand the impact of the clustering on the actual cancer progression inference, we considered SCITE [17], SASC [5] and SPhyR [7], three published, publicly available inference tools tailored to SCS data, and have been shown to scale to instances of the size we consider in our study — the latter tool does clustering using *k*-means as part of the inference. We executed SCITE [17] and SPhyR [7] on both the obtained clusters and on the unclustered datasets, to understand the effect of the clustering on the tools. SASC was not able to complete in a reasonable amount of time on the unclustered data, due to the higher complexity of the search space of the solutions it generates. We performed the inference with all of the clustering methods used as a preprocessing step. Note that all methods were run with default parameters, with the exception of SASC and SPhyR being parameterized to output trees with no losses, in order to be able to compare with SCITE, which does not model losses, i.e., rather, it adheres to the infinite sites assumption.

For assessing the accuracy of the methods we used the measures defined in [5], [6]:

**Ancestor-descendant accuracy**: for each pair of mutations in an ancestor-descendant relationship in the ground truth, we check whether the relationship is conserved in the inferred tree (*TP*) or whether it is not (*FN*). For each pair of mutations in an ancestor-descendant relationship in the inferred tree, we also check if such relationship does not exist in the ground truth tree (*FP*).
**Different lineages accuracy**: similarly to the previous measure, we check whether mutations in different branches are correctly inferred or if any pair of mutation is erroneously inferred in different branches.

Figures 11, 12 and 13 show very interesting results. Regarding the results of SCITE and SASC, as expected, we observe a severe drop in performance when used in combination with *k*-means, affinity, agglomerative, BIRCH and spectral clustering methods. This fact is supported by the gap in the precision of the methods — a low precision indeed leads to a low accuracy in the tree reconstruction. The trend is still present in the different lineages accuracy, but to a lesser extent — this is because, as previously discussed, a cancer inference method can separate *clusters* of mutations, but when a cluster is computed it is not possible to separate mutations *within* this cluster.

**Fig. 11.**
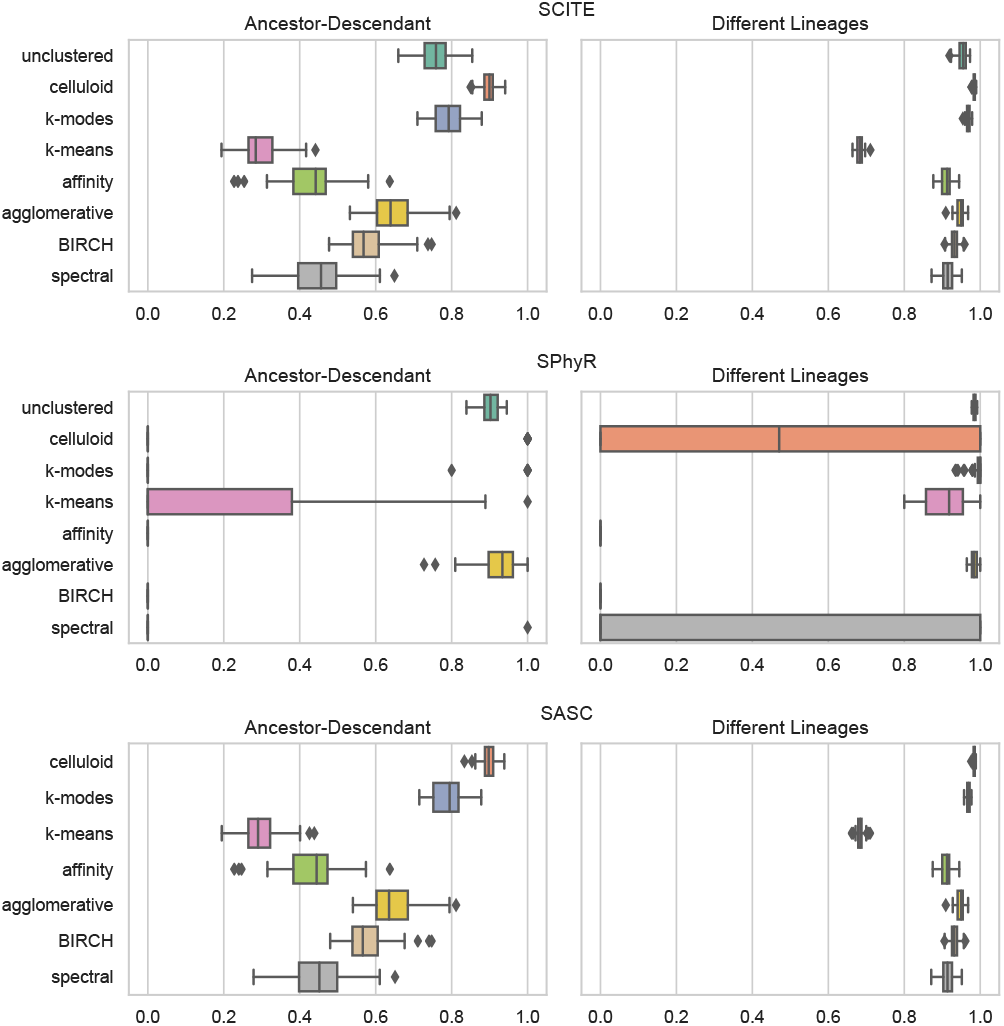
Ancestor-descendant and different lineages accuracy measures for experiment 1. Cancer phylogenies are inferred by SCITE (Top), SPhyR (Middle) and SASC (Bottom) using as input both the unclustered data as well as the clusters obtained by *celluloid*, ***k***-modes, ***k***-means, affinity, agglomerative, BIRCH and spectral clustering.

**Fig. 12.**
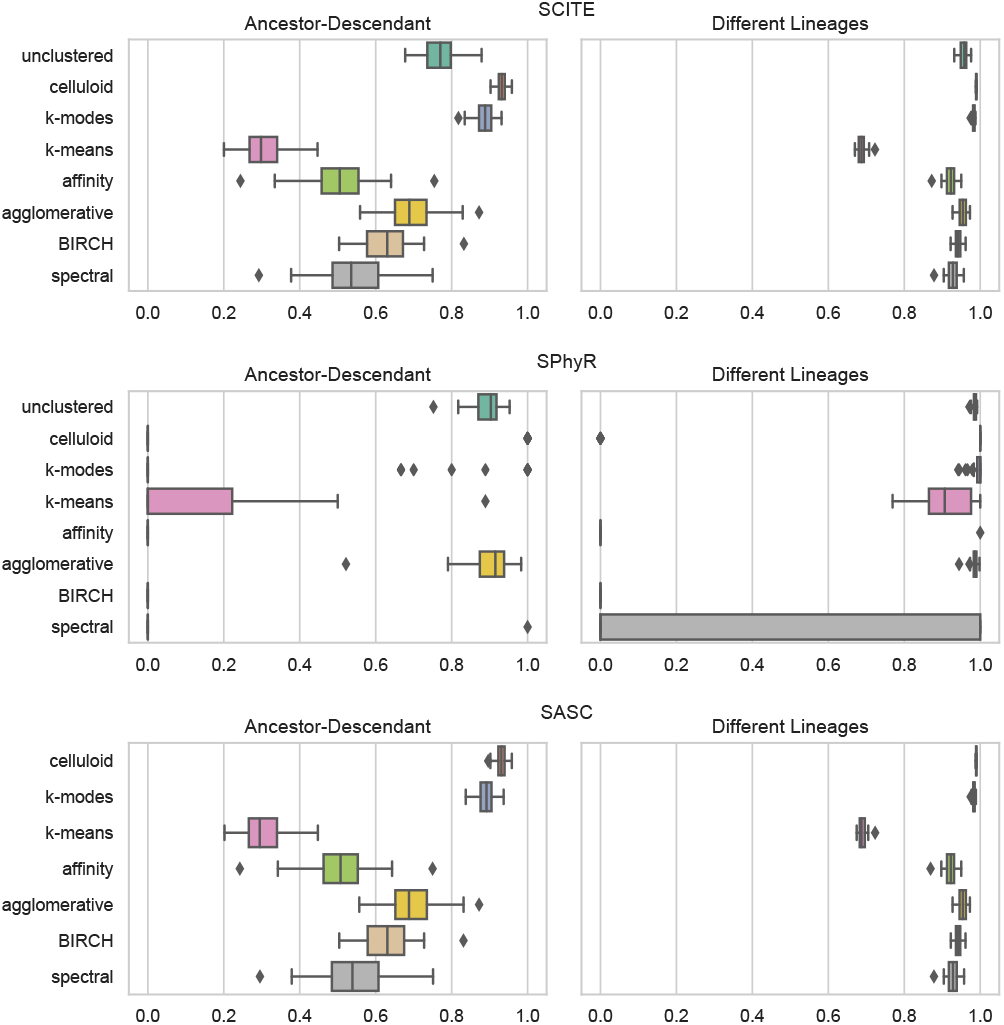
Ancestor-descendant and different lineages accuracy measures for experiment 2. Cancer phylogenies are inferred by SCITE (Top), SPhyR (Middle) and SASC (Bottom) using as input both the unclustered data as well as the clusters obtained by *celluloid*, ***k***-modes, ***k***-means, affinity, agglomerative, BIRCH and spectral clustering.

**Fig. 13.**
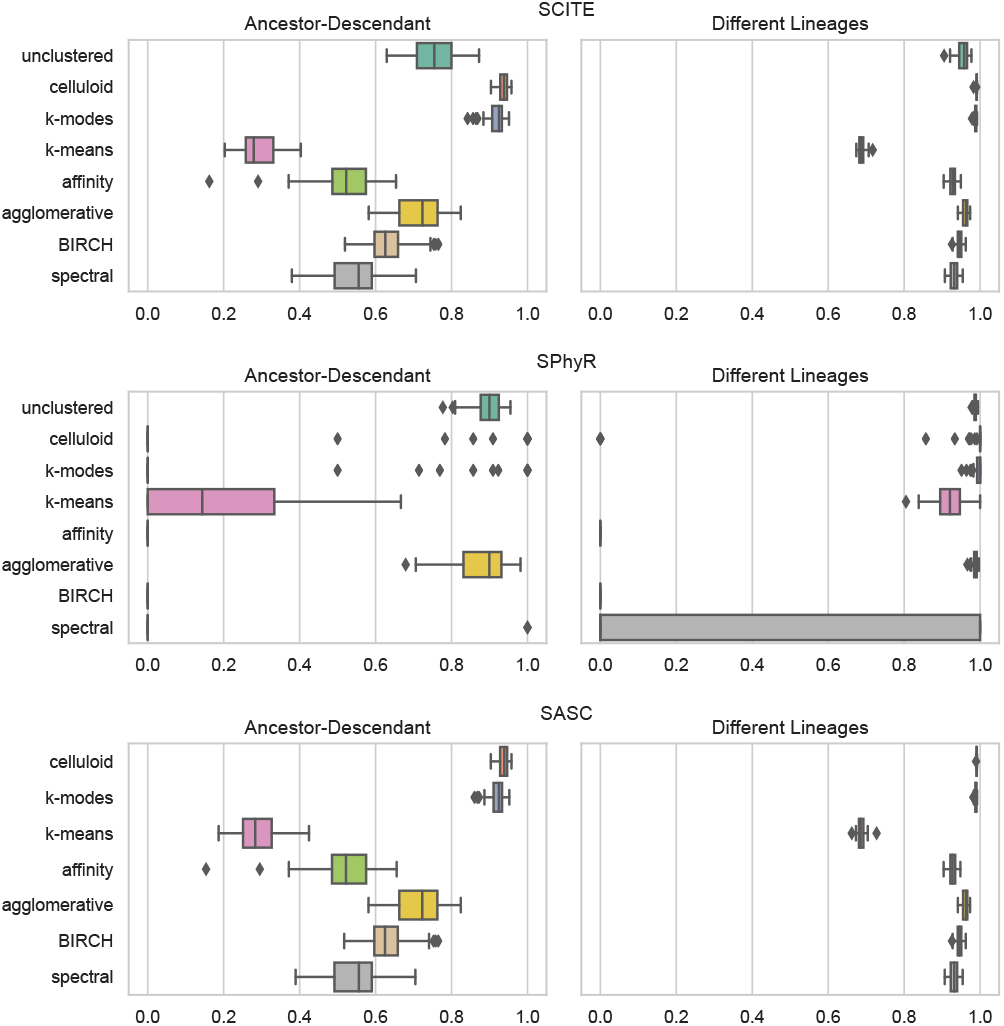
Ancestor-descendant and different lineages accuracy measures for experiment 3. Cancer phylogenies are inferred by SCITE (Top), SPhyR (Middle) and SASC (Bottom) using as input both the unclustered data as well as the clusters obtained by *celluloid*, ***k***-modes, ***k***-means, affinity, agglomerative, BIRCH and spectral clustering.

On the other hand, the results obtained by SCITE when *celluloid* and *k*-modes are used are much better, in particular, *celluloid* as a preprocessing step — which leverages our conflict dissimilarity — allows SCITE to score higher in both ancestor-descendant and different lineages accuracies than it is able to using unclustered data as input. For this inference tool, using *celluloid* as a preprocessing step actually helps SCITE to achieve a better score than without it — experiment 3, being the largest of the three, shows the biggest improvements. SCITE on unclustered data scores an average of 0.7551 and 0.9537 for ancestor-descendant and different lineages respectively, against an average of 0.9358 and 0.9907 after clustering the datasets using *celluloid*. Moreover, *celluloid* allows SCITE to achieve a 20x speedup in runtime, on average.

The results obtained by SASC, shown in Figures 11, 12 and 13, are very similar to what is computed by SCITE — in particular, *celluloid* provides much better results than any other clustering method. Once again, on the larger experiment, the gap between the clustering methods is seen the most — in particular, SASC scores an average of 0.9365 and 0.9909 for ancestor-descendant and different lineages respectively, when *celluloid* is used as a preprocessing step. These results are particularly interesting since SASC is unable to complete in a reasonable amount of time on the unclustered data, hence the reason for this absence from the experiments.Unlike the previous tools, SPhyR shows very interesting results since the method itself performs a *k*-means clustering on both mutations and cells given as input. For this reason, clustering already clustered data can be a hit-or-miss case, as seen in Figures 11, 12 and 13. In almost all cases, preprocessing the data causes SPhyR to obtain extremely low values in the ancestor-descendant measure, while achieving a very good different lineages score, with the exception of spectral clustering in all experiments and *celluloid* in experiment 1. On the other hand, the agglomerative clustering seems to have no impact or to slightly improve the tool — this could be a consequence of the high recall and low precision achieved by the clustering algorithm — indeed it is possible that separating well clusters but not merging them very much leads the clustering step of SPhyR to achieve a better result.

### C. Application on real data

Finally, we run the entire pipeline (clustering + inference method) on an oligodendroglioma IDH-mutated tumor [36] consisting of 1842 mutations over 926 cells, to assess if the improvement in the runtimes that we have seen on simulated data carry over to a real dataset.

Such computation was performed using SCITE, which on unclustered data, takes as long as 68 hours, while adding a preprocessing step with *celluloid*, we were able to compute the tree within 1 hour, therefore decreasing the time needed by a factor of 70. Moreover, it is most likely that preprocessing the data with *celluloid* provides a better phylogeny than SCITE alone, as seen in the experiments on simulated data.

Furthermore we computed the same dataset using SASC, for which such computation was previously not possible with the use of SASC alone, due to the running time required — SASC was not able to provide a solution in a time-frame of three weeks. Figure 14 shows the tree inferred, where each node is one of 100 clusters computed by *celluloid*.

**Fig. 14.**
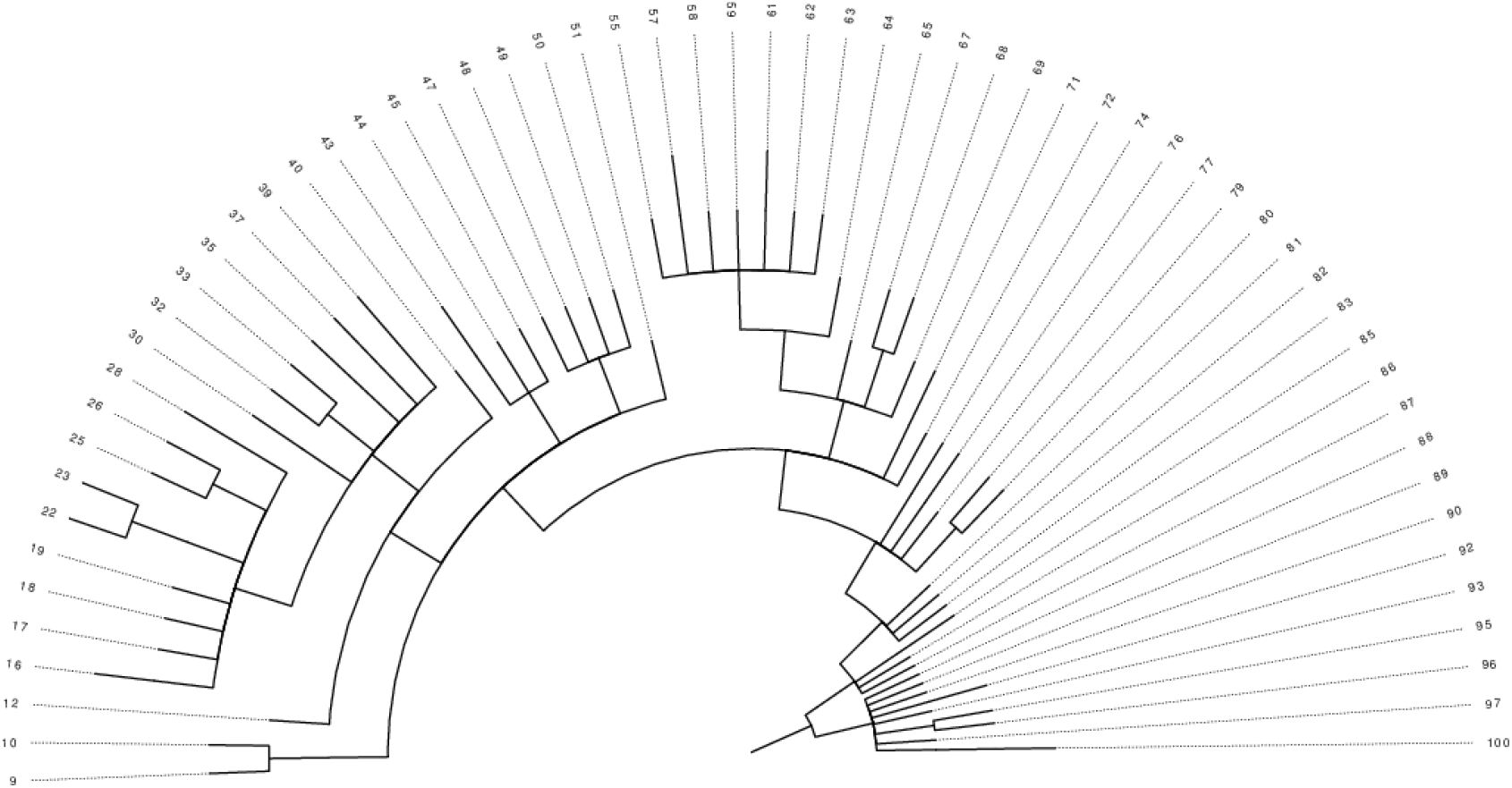
Tree computed on the oligodendroglioma IDH-mutated tumor from [36]. The labels on the nodes represent the 100 clusters obtained by *celluloid*. The tree was inferred using the cancer inference tool SASC.

## IV. Format interchange

Since *celluloid* is designed to be a part of any pipeline that infers a tumor phylogeny, we need to be able to produce data that can be fed as input to a variety of tools. For this reason, we have implemented a feature that translates between the file formats used by different programs: SASC, SCITE, and SPhyR. To allow future extensions, we have designed this part around a *Parsing Expression Grammar* (PEG) [41]; a family of context-free grammars that admit deterministic parsers.

PEGs are syntactically similar to context-free grammars (CFGs) but present different interpretations; in particular the choice operator is prioritized, meaning that they will test alternative models in order, using the first successful match. For example, while the rules *A* → *ab*|*a* and *A* → *a*|*ab* are equivalent in a CFG, those rules in a PEG have a different semantics, *e.g.*, the second choice of the second rule will never succeed if the input string starts with *a*. This restriction allows to have faster parsers.

For obtaining a parser for each PEG, we used TatSu [51] which is an open source tool that starts from a grammar (in our case a PEG) and produces a *Packrat parser* [42]. Packrat parsers guarantee a linear parse time, at the possible expense of a large memory usage. In our case, this is not a problem since the grammars involved are quite simple.

With *celluloid*, we provide commands to translate between the inputs of the inference methods present in this paper, as well as their grammars. It is easy to add new custom grammars to the translator, and so, as new methods — which use custom formats — are developed, they can be easily and seamlessly incorporated into this translation framework.

## V. Discussion

Motivated by the advancements in single cell sequencing (SCS) technologies such as decreasing costs and improved quality which will result in larger SCS instances, we proposed a method *celluloid* to reduce the size of such instances, via categorical clustering based on the *k*-modes framework, and a novel *conflict dissimilarity* — tailored to properties specific to SCS instances. The idea is that this can allow tools which infer cancer phylogenies purely from SCS data, such as SCITE [17], SPhyR [7] or SASC [5] — which work well on the instances of today — to scale to the size of SCS instances of the near future. We hence devised *celluloid* for clustering purely single-cell data, and focused on comparing other methods with this same goal in mind. Notwithstanding, there are important *hybrid* methods for inferring cancer phylogenies from a combination of bulk sequencing and SCS data [24], [33], which even perform clustering as part of the phylogeny inference (like with the purely SCS tool, SPhyR).

We have shown how to compare various clustering methods on single cell sequencing data, with three distinct experiments. More precisely, we describe two experiments on synthetic data: the first experiment measures the quality of the clusters computed by the methods, while the second experiment considers the effect of the clustering on the quality of trees obtained downstream from a phylogeny inference tool. Finally, we have described an experiment on real data that measures the usefulness of the clustering procedure, by computing the improvement in the runtime for a large instance.

We have made available the entire pipeline [43] that runs the clustering tools on those data, and computes the plots and tables used in the cluster analysis. Our comparison is reproducible and can be easily extended by modifying a Snakemake [21] file.

One of the main conclusions that we can draw from our comparison is that *k*-means is not an adequate choice for clustering single cell data for inferring tumor phylogenies. The second main finding of our paper is that a suitable clustering step, such as *celluloid*, not only decreases the runtime of the phylogeny inference methods, but can also *improve* the quality of the inferred phylogenies, as we have shown for SCITE and SASC.

Future work includes a more complete and comprehensive study of clustering methods and downstream cancer phylogeny inference methods, but also of simulated and real datasets.

Since the *k*-modes framework saw a boost in performance when coupled with our novel conflict dissimilarity, an interesting future work is to determine if there are other dissimilarities, or even other categorical clustering frameworks which perform even better. For example, the conflict dissimilarity takes into account the high drop-out rate in SCS data; maybe there are refinements that could account for other aspects of SCS data such as heterogeneous false-negative rate [5]. More generally, maybe there are other dimensionality reduction techniques, or ways to select or extract a compact set of representative features of this dataset (not necessarily mutations or cells), which could in turn be passed to some downstream phylogeny inference.

Finally, while we focused on single cell data in this study, a more long-term research direction is to develop tools — which use some combination of clustering + phylogeny inference — for inferring cancer phylogenies using multiple sources of information together (*e.g*., transcriptomics data, health informatics data, phylogenetic information of other closely-related cancers, *etc*.).

## Acknowledgment

We thank Iman Hajirasouliha and Dana Silverbush for several illuminating discussions. We also thank an anonymous reviewer for pointing out a possible problem in our initial model formulation. We also thank Danilo Fumagalli for having implemented the format interchange procedure.

