## Supplementary Material for "Effective Clustering for Single Cell Sequencing Cancer Data"

May 14, 2021

The following figures depict results for experiments on simulated data (described in Section II-H of the main text) for parameter settings supplementary to those of the main text.

Figs. 1–9 are analogous to those of Figs. 2–10 of the main text but fix the number of cells to 300, while the number of mutations varies from 300, 500 to 700. While such number(s) of mutations is small enough to be directly input to current cancer phylogeny inference methods (also the reason for the choice of a smaller clustering size of $k = 50$), clustering results nonetheless reveal similar trends as the experiments of the main text.

Figs. 10 and 11 consider the same number (1000) of mutations as those of Figs. 2–4 of the main text, but consider larger numbers, 400 and 500 of cells, respectively. Here we see that *celluloid* still dominates in terms of precision, while recall tends to plateau for all methods.

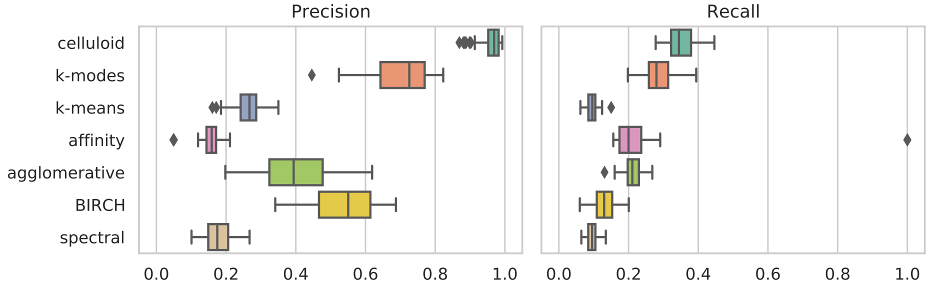


Figure 1: Precision and recall results, generated with a total of **300** mutations, 300 cells and a clustering size of $k = 50$. The plots include results for *celluloid*, $k$-modes, $k$-means, affinity, agglomerative, BIRCH and spectral clustering.

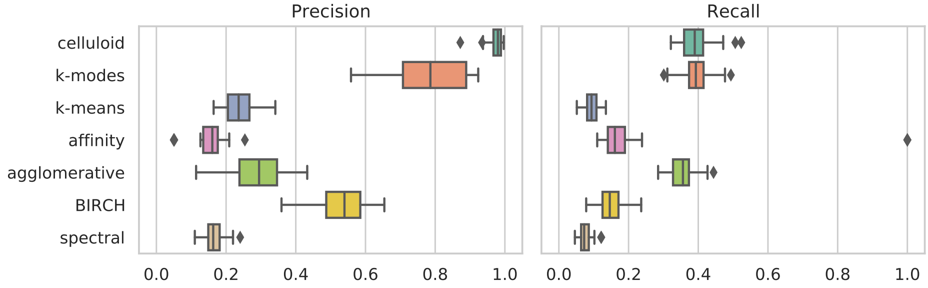


Figure 2: Precision and recall results, generated with a total of **500** mutations, 300 cells and a clustering size of $k = 50$. The plots include results for *celluloid*, $k$-modes, $k$-means, affinity, agglomerative, BIRCH and spectral clustering.

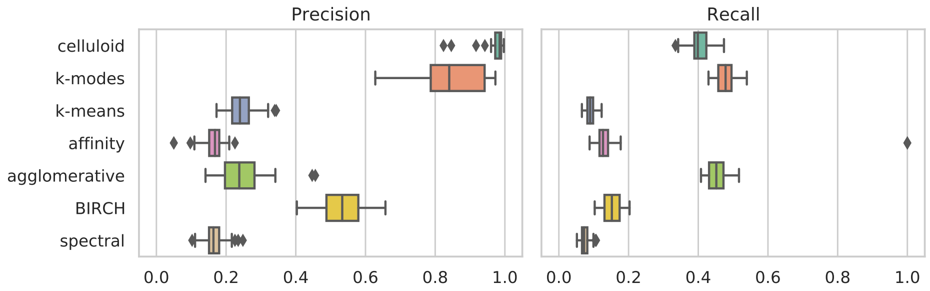


Figure 3: Precision and recall results, generated with a total of **700** mutations, 300 cells and a clustering size of $k = 50$. The plots include results for *celluloid*, $k$-modes, $k$-means, affinity, agglomerative, BIRCH and spectral clustering.

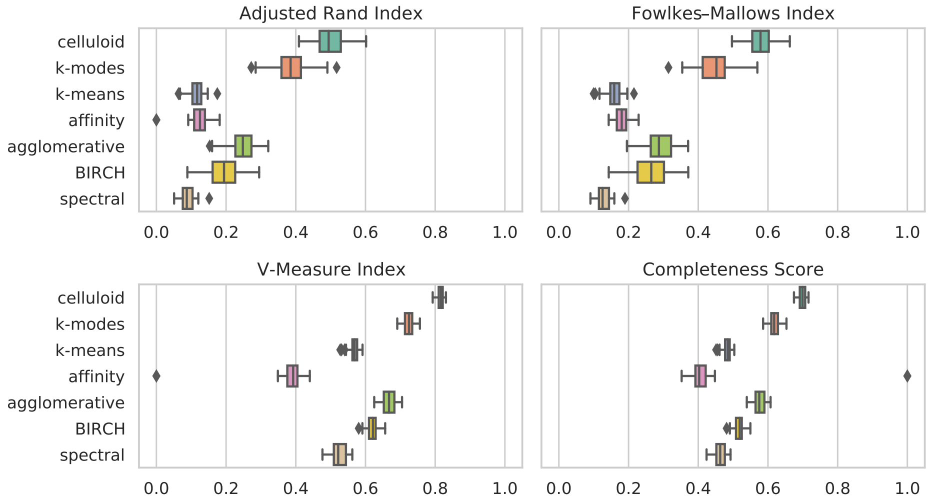


Figure 4: The adjusted Rand index, Fowlkes-Mallows index, completeness score and V-measure between all clustering methods and the ground truth generated with a total of **300** mutations, 300 cells and a clustering size of $k = 50$. The plots include results for *celluloid*, $k$-modes, $k$-means, affinity, agglomerative, BIRCH and spectral clustering.

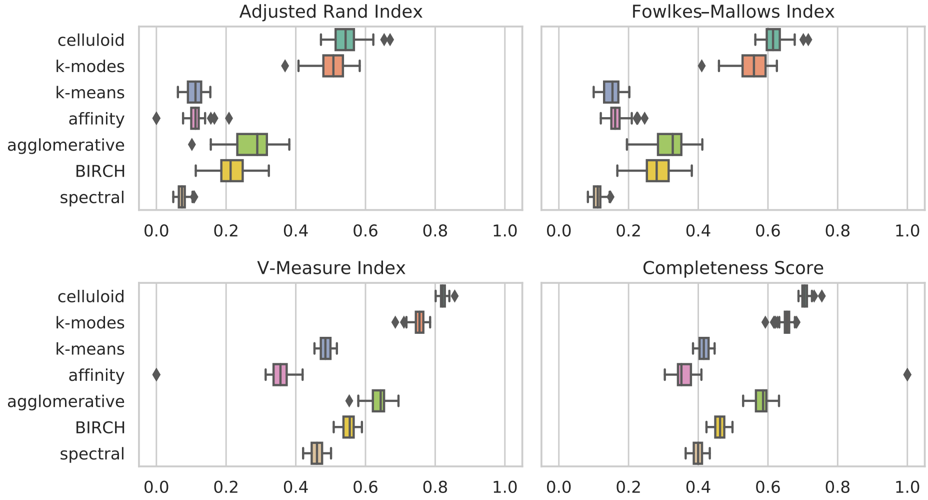


Figure 5: The adjusted Rand index, Fowlkes-Mallows index, completeness score and V-measure between all clustering methods and the ground truth generated with a total of **500** mutations, 300 cells and a clustering size of $k = 50$. The plots include results for *celluloid*, $k$-modes, $k$-means, affinity, agglomerative, BIRCH and spectral clustering.

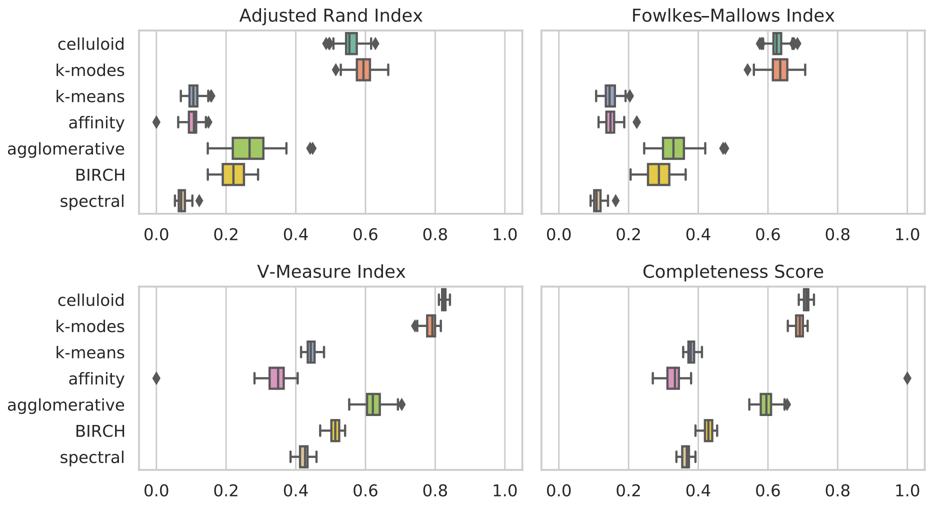


Figure 6: The adjusted Rand index, Fowlkes-Mallows index, completeness score and V-measure between all clustering methods and the ground truth generated with a total of **700** mutations, 300 cells and a clustering size of $k = 50$. The plots include results for *celluloid*, $k$-modes, $k$-means, affinity, agglomerative, BIRCH and spectral clustering.

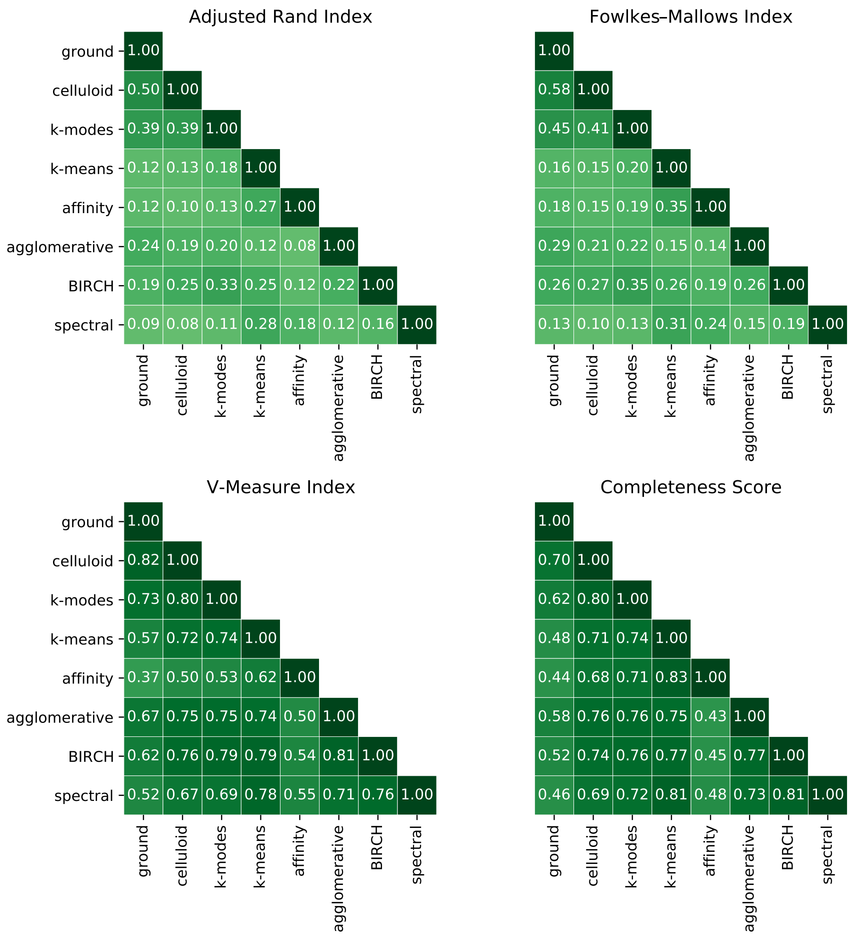


Figure 7: The adjusted Rand index, Fowlkes-Mallows index, completeness score and V-measure between all pairs of clustering methods generated with a total of **300** mutations, 300 cells and a clustering size of $k = 50$. The plots include results for *celluloid*, $k$-modes, $k$-means, affinity, agglomerative, BIRCH and spectral clustering. Each cell is the average of the scores obtained for each simulated instance.

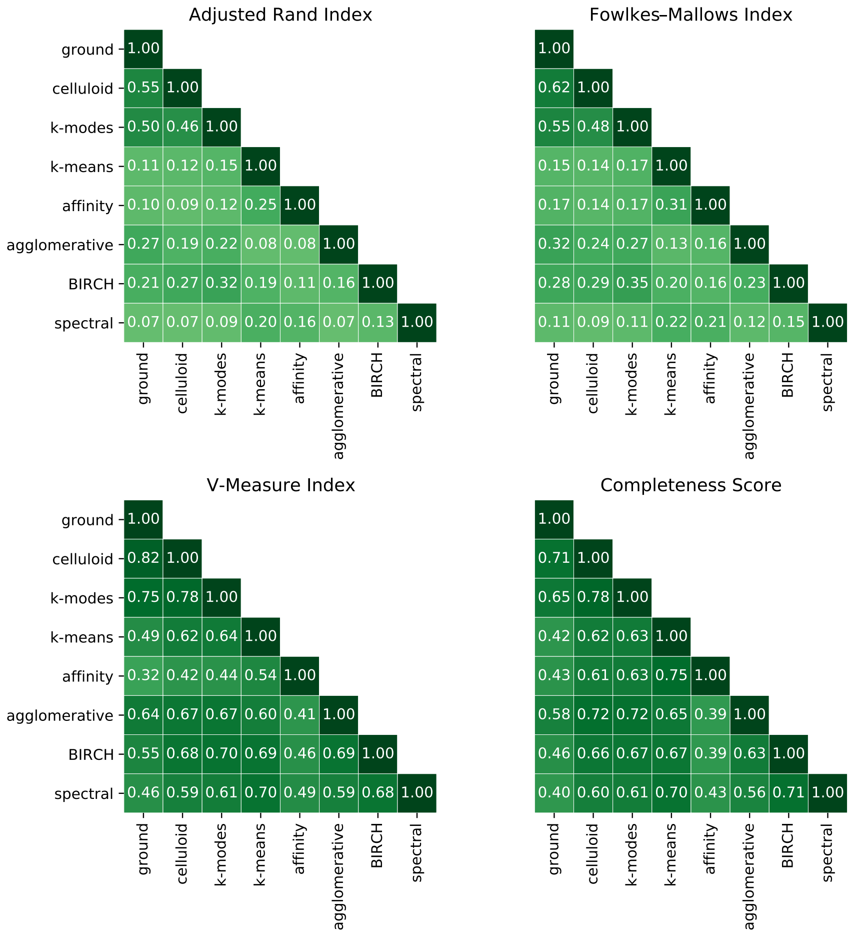


Figure 8: The adjusted Rand index, Fowlkes-Mallows index, completeness score and V-measure between all pairs of clustering methods generated with a total of **500** mutations, 300 cells and a clustering size of $k = 50$. The plots include results for *celluloid*, $k$-modes, $k$-means, affinity, agglomerative, BIRCH and spectral clustering. Each cell is the average of the scores obtained for each simulated instance.

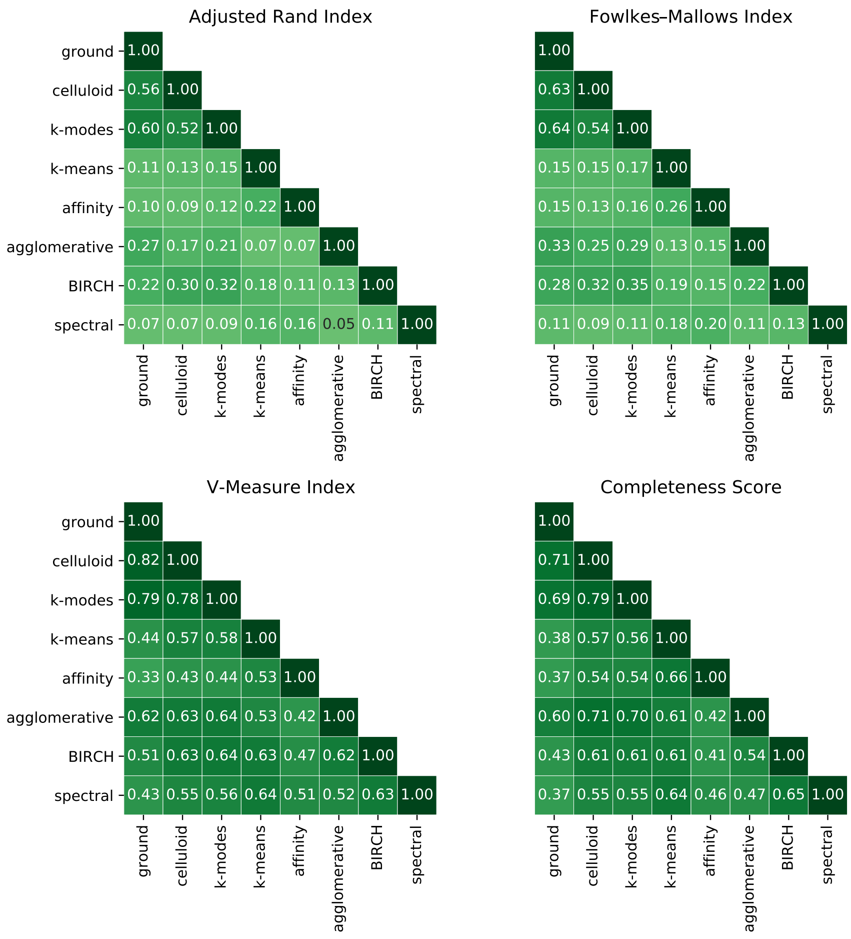


Figure 9: The adjusted Rand index, Fowlkes-Mallows index, completeness score and V-measure between all pairs of clustering methods generated with a total of **700** mutations, 300 cells and a clustering size of $k = 50$. The plots include results for *celluloid*, $k$-modes, $k$-means, affinity, agglomerative, BIRCH and spectral clustering. Each cell is the average of the scores obtained for each simulated instance.

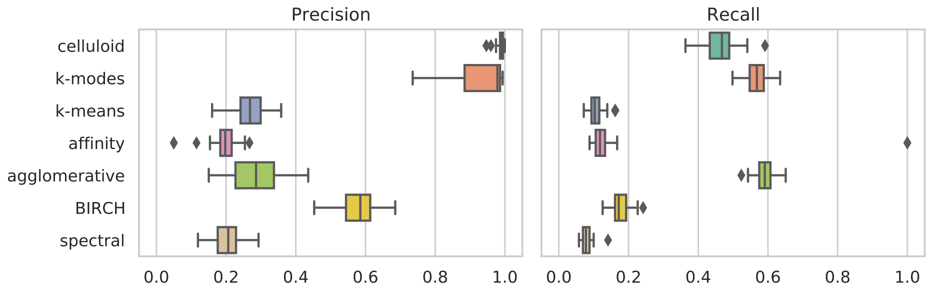


Figure 10: Precision and recall results, generated with a total of 1000 mutations, **400** cells and a clustering size of k = 100. The plots include results for celluloid, k-modes, k-means, affinity, agglomerative, BIRCH and spectral clustering.

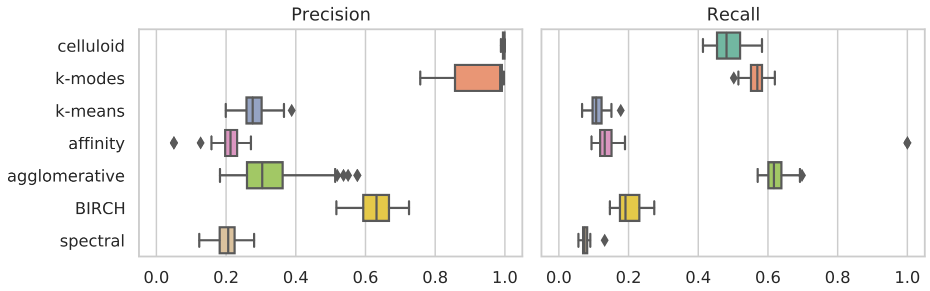


Figure 11: Precision and recall result, generated with a total of 1000 mutations, **500** cells and a clustering size of k = 100. The plots include results for celluloid, k-modes, k-means, affinity, agglomerative, BIRCH and spectral clustering.
